# *Mycobacterium tuberculosis* infection elevates SLIT2 expression to modulate oxidative stress responses in macrophages

**DOI:** 10.1101/2022.10.13.512188

**Authors:** Salik Miskat Borbora, Sneha Bhatt, Kithiganahalli Narayanaswamy Balaji

**Author notes:** To whom correspondence should be addressed: Dr. Kithiganahalli Narayanaswamy Balaji, Department of Microbiology and Cell Biology, Indian Institute of Science, Bangalore 560012, Karnataka, India.

## Abstract

Mycobacterium tuberculosis (Mtb), the causative agent of the pulmonary ailment, tuberculosis (TB), continues to thrive owing to a disorganized immune response against it by the host. Among other factors, the rewiring of distinct host signaling pathways is effectuated by the intracellular bacterium, resulting in pathogen-favorable outcomes. Oxidative stress build-up is a key cellular manifestation that occurs during mycobacterial infection. Enhanced oxidative stress is brought about by the cumulative effect of elevated reactive oxygen species generation as well as the inept ability of the cell to mitigate ROS levels. Here, we report the increased expression of the neuronal ligand, SLIT2, during mycobacterial infection in macrophages. By employing loss of function analysis using specific inhibitors, we attribute the heightened expression of SLIT2 to the Mtb-mediated phosphorylation of the p38/JNK pathways. Also, using chromatin immunoprecipitation (ChIP) analysis, we found reduced levels of the repressive H3K27me3 signature on the Slit2 promoter during mycobacterial infection. Furthermore, SLIT2 was found to promote the expression of cellular pantetheinase, Vanin1 (VNN1), that contributed to copious levels of ROS within the macrophage cellular milieu. Thus, we dissect essential molecular details leading to the robust expression of SLIT2 during Mtb infection while outlining the potential consequences of SLIT2 upregulation in infected macrophages.

**Graphical abstract:** 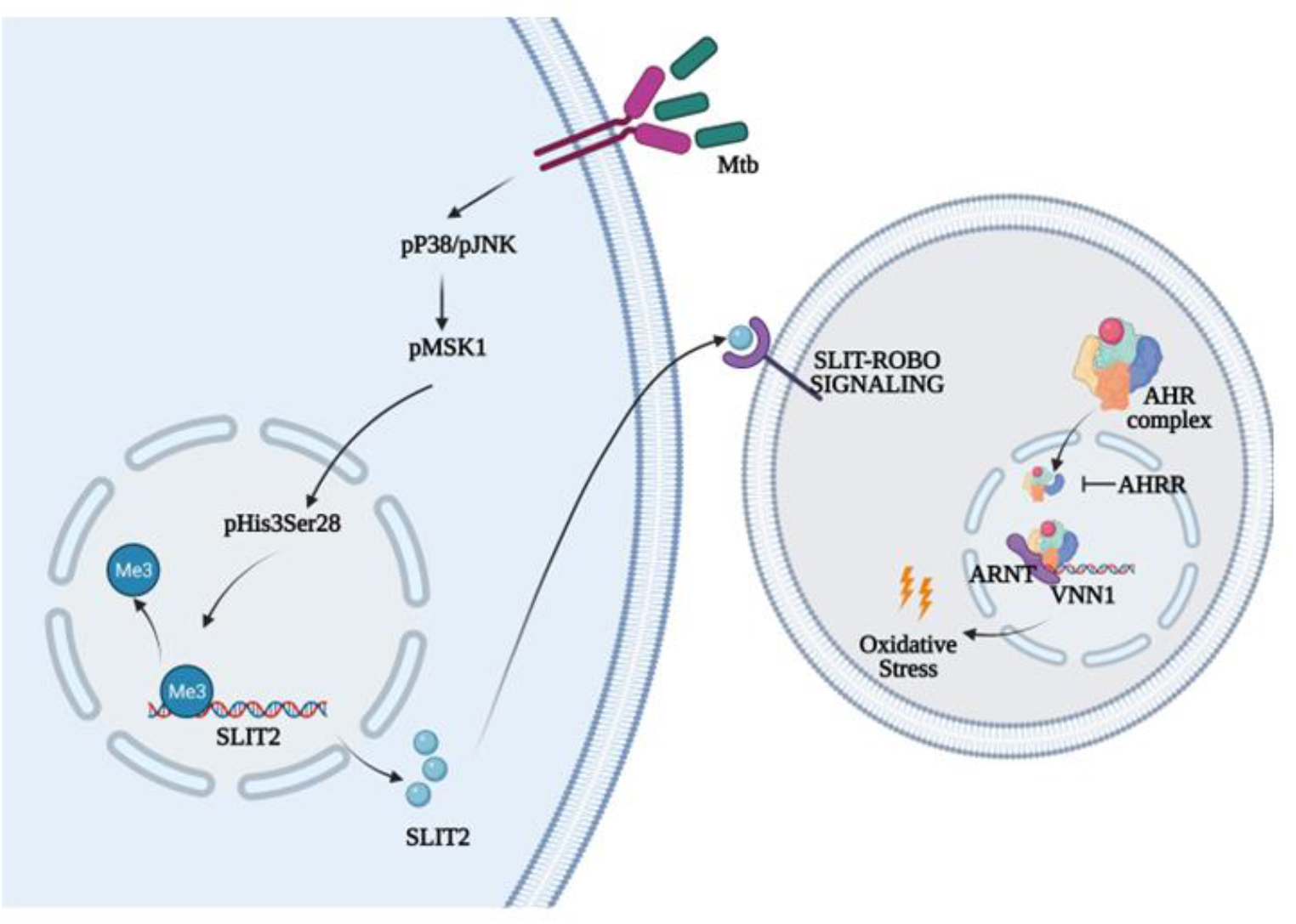

## Introduction

The intracellular microbe, *Mycobacterium tuberculosis* (Mtb) triggers a series of changes within the host that enhances the scope of pathogen survival and dissemination. Macrophages are one of the primary cells infected by Mtb wherein anti-mycobacterial processes are initiated alongside a focused inflammatory response(1). Available reports have implicated the reprogramming of cellular signaling pathways as key events in the establishment and perpetuation of TB infection(2). Notably, diverse signaling pathways viz. NOTCH(3, 4), WNT(5), Sonic Hedgehog(6) etc. have been reported to be differentially regulated upon mycobacterial infection. Besides, Ca^2+^ signaling(7), phosphatidylinositol-4,5-bisphosphate 3-kinase (PI3K)(8), and Janus Kinase/ Signal transducer and activator of transcription (JAK-STAT)(9, 10), have also been observed to be altered during Mtb infection.

SLIT and ROBO (ROUNDABOUT) proteins constitute a distinctive signaling pathway that was first reported to mediate axon repulsion in the developing nervous system(11). SLITs are secreted glycoproteins, while ROBO receptors belong to the immunoglobulin superfamily of cell adhesion molecules (CAMs). In mammals, there are three SLIT ligands and four Robo receptors(12). Although, initial reports indicated the presence of these molecules in neural tissues, subsequently they have been observed across the mammalian system(13). Concurrently, SLIT proteins have been associated with non-neuronal cellular events such as angiogenesis(14, 15), leukocyte chemotaxis(16), fibrosis(17), ROS generation(18) etc. Further, SLIT2 has been shown to be responsive during LPS-driven inflammatory responses in endothelial cells(19). To our interest, such cellular events have also been reported during TB progression(20, 21). Thus, it is intriguing to explore the role of SLIT proteins in Mtb-mediated changes within infected macrophages.

During Mtb infection, macrophages encounter oxidative stress that stems from an uneven balance between the oxidant and antioxidant machinery of the cell(22). Within the macrophages, Mtb employs its ROS scavenging enzyme KatG, cell surface antioxidants like alpha-keto acid dehydrogenase, mycothiol etc. to overcome the hostile environment(23, 24). Unchecked ROS levels would be detrimental in any cell as it aids lipid oxidation and peroxidation, culminating in cell death and tissue destruction(25, 26).

In this context, it is imperative to limit ROS within infected macrophages to prevent pathogen dissemination. A gamut of antioxidant enzymes such as HO1, TRXR1 have been implicated in maintaining redox homeostasis during Mtb infection(27, 28). The tripeptide, glutathione is yet another molecule that has been shown to protect cells from oxidative damage(29). With regards to TB, glutathione has also been demonstrated to modulate immune responses wherein supplementation of NAC (N-acetyl cysteine), a glutathione precursor, could inhibit Mtb growth in blood cultures(30). However, there are scant reports on specific proteins that cripple the host cell response upon ROS build-up during Mtb infection.

Here, we identify an epigenetic basis of SLIT2 upregulation during Mtb infection. Concurrently, we delineate the molecular details about SLIT2-mediated ROS accumulation in Mtb-infected macrophages. By using specific inhibitors, we establish VNN1, a cellular pantetheinase, as a key molecule in regulating oxidative stress responses during Mtb infection. Furthermore, *in vitro* CFU assay revealed compromised mycobacterial survival upon VNN1 inhibition.

## Results and Discussion

### Mtb-mediated elevated SLIT2 expression contributes to oxidative stress in macrophages

To determine the role of SLIT-ROBO signaling pathway during Mtb infection, we analyzed the relative expression of SLIT ligands in both *in vitro* and *in vivo* scenarios. Of the three ligands, murine macrophages elicited robust expression of SLIT2 at the transcript and protein level **(Fig.1A,B)**. Similarly, mycobacterial infection resulted in enhanced expression of SLIT2 in THP1 macrophages **(Fig.S1A)**. These results were corroborated in our *in vivo* experiments as well wherein elevated SLIT2 transcripts and protein were observed in the lung homogenates of mice infected with Mtb H37Rv **(Fig.1C,D)**. Likewise, assessment of the status of ROBO receptors revealed enhanced expression of the ROBO2 receptor in peritoneal macrophages **(Fig.S1B,C)**. In neuronal cells, SLIT2 has been extensively implicated in axon guidance that contribute to synaptic connections(31). Besides, studies have implicated the role of Reactive oxygen species in neuronal processes such as growth cone guidance, neuronal migration, neuronal polarity etc.(32, 33). Again, Mtb infection has been associated with changes in ROS levels in macrophages(22). Thus, we surmised a possible correlation between SLIT2 production and ROS levels in macrophages upon Mtb infection. To our interest, a recent study delineated the role of SLIT2-mediated growth cone guidance in a ROS-dependent manner(18). Immunofluorescence studies using CellROX Deep Red reagent revealed compromised ROS levels in murine macrophages wherein SLIT2 levels were curbed using specific siRNA **(Fig.1E,F)**. Taken together, Mtb-mediated SLIT2 expression augmented ROS levels in macrophages.

**Figure 1.**
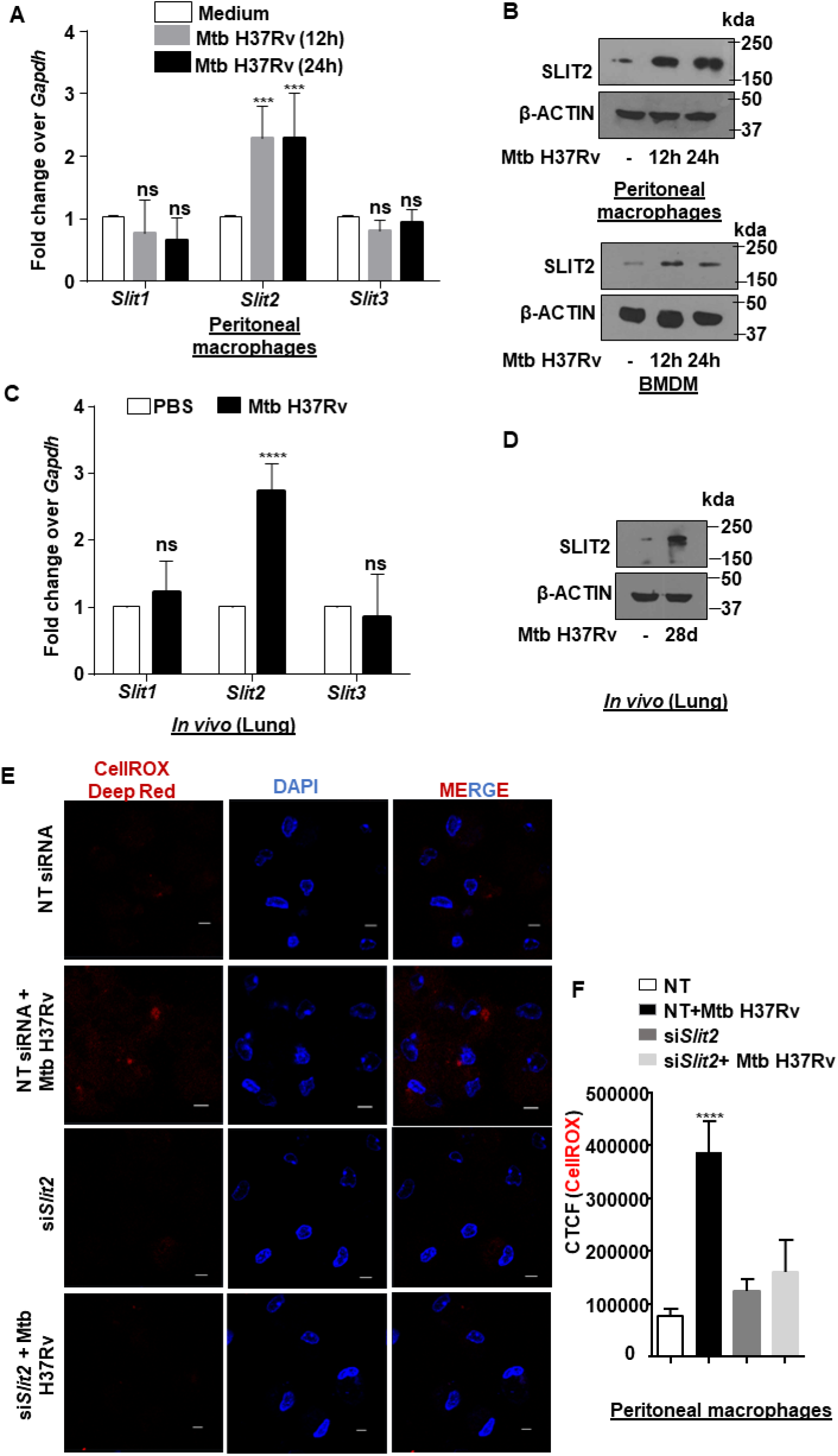
Mtb infection increases SLIT2 expression in macrophages that enhances ROS levels. **(A)** Mouse peritoneal macrophages were infected with Mtb H37Rv for the indicated time points and assessed for the transcript levels of SLIT ligands. **(B)** Mouse peritoneal macrophages *(top)* and bone marrow-derived macrophages (BMDM) *(below)* were infected with Mtb H37Rv for the indicated time points. Whole cell lysates were assessed for SLIT2 protein expression by immunoblotting. **(C)** BALB/c mice were aerosol-infected (~100CFU) with Mtb H37Rv for 28 days. Transcript levels of SLIT ligands was analyzed in the lung homogenates of uninfected and infected mice by qRT-PCR**. (D)** BALB/c mice were aerosol-infected with Mtb H37Rv for 28 days. Protein levels of SLIT2 was analyzed in the lung homogenates of uninfected and infected mice by immunoblotting. **(E, F)** Mouse peritoneal macrophages were transfected with NT or *Slit2* siRNAs. Transfected cells were infected with Mtb H37Rv for 24 h and assessed for the accumulation of ROS by fluorescence microscopy (CellROX Deep Red); **(E)** representative image and **(F)** respective quantification. All immunoblotting and immunofluorescence data are representative of three independent experiments. Lung homogenates from at least three mice/ group (PBS treated and Mtb H37Rv infected) were independently assessed for SLIT ligands expression by qRT-PCR and SLIT2 expression by immunoblotting. β-ACTIN was utilized as loading control. qRT-PCR data represents mean±S.E.M. from three independent experiments. ROS, reactive oxygen species; h, hours; d, days; kda, kilodalton; CTCF, corrected total cell fluorescence. *** p<0.001; ****, p < 0.0001 (Student’s t-test in A,C; One way ANOVA in F; GraphPad Prism 6.0 and 9.0). Scale bar, 5 μm.

### SLIT2 expression in macrophages is epigenetically regulated during Mtb infection

Available studies indicated an epigenetic basis of SLIT2 regulation, among others. Notably, SLIT2 promoter is hypermethylated in various cancerous scenarios(34–37), while specifically it has been shown to be a target of repression by EZH2 in prostate cancer cases(38). The histone-lysine N-methyltransferase enzyme, EZH2, has been shown to regulate gene expression in inflammatory disease conditions(39, 40). Thus, we hypothesized that in the uninfected scenario, SLIT2 expression is suppressed due to enhanced H3K27me3 mark over its promoter. However, upon Mtb infection, dynamic epigenetic changes result in reduction of the H3K27me3 mark and consequent elevation of *Slit2* expression **(Fig.2A)**. Chromatin immunoprecipitation (ChIP) assay in THP1 macrophages revealed decreased H3K27me3 signature on *Slit2* promoter upon Mtb infection **(Fig.S2A)**. To our interest, we came across reports which suggested that stress-responsive kinases (p38/JNK/MEK) could mediate changes in the epigenetic landscape of genes, thereby contributing to the expression or repression of genes(41). Notably, phosphorylation of the Serine 28 residue of Histone3 (H3Ser28) leads to enhanced demethylation or acetylation at H3K27 residue, leading to transcription initiation(42). Mtb-infected macrophages displayed enhanced phosphorylation of Histone3 (H3Ser28) **(Fig.S2B,C).** A ChIP assay revealed the presence of phosphorylated H3Ser28 on *Slit2* promoter with a concomitant decrease in the occupancy of H3K27me3 signature **(Fig.2B)**. Phosphorylated JNK1/2 and/or p38 can phosphorylate MSK1 (43), which is pivotal for H3Ser28 phosphorylation. Perturbation of p38, JNK1/2, and MSK1 phosphorylation using specific inhibitors (SB203580 for p38; SP600125 for JNK1/2; H-89 for MSK1) displayed compromised expression of SLIT2 upon Mtb infection at the transcript level **(Fig.2C)** and protein level **(Fig.2D)**. Thus, we identify p38, JNK1/2 and MSK1 as important upstream regulators of SLIT2 expression during Mtb infection. A parallel investigation has shown that JNK signaling is central to the activation of the SLIT-ROBO pathway in epithelial cells during wound repair(44). Subsequently, inhibition of p38 and JNK1/2 phosphorylation compromised ROS levels upon Mtb infection, corroborating our initial observations with SLIT2 **(Fig.2E,F)**.

**Figure 2.**
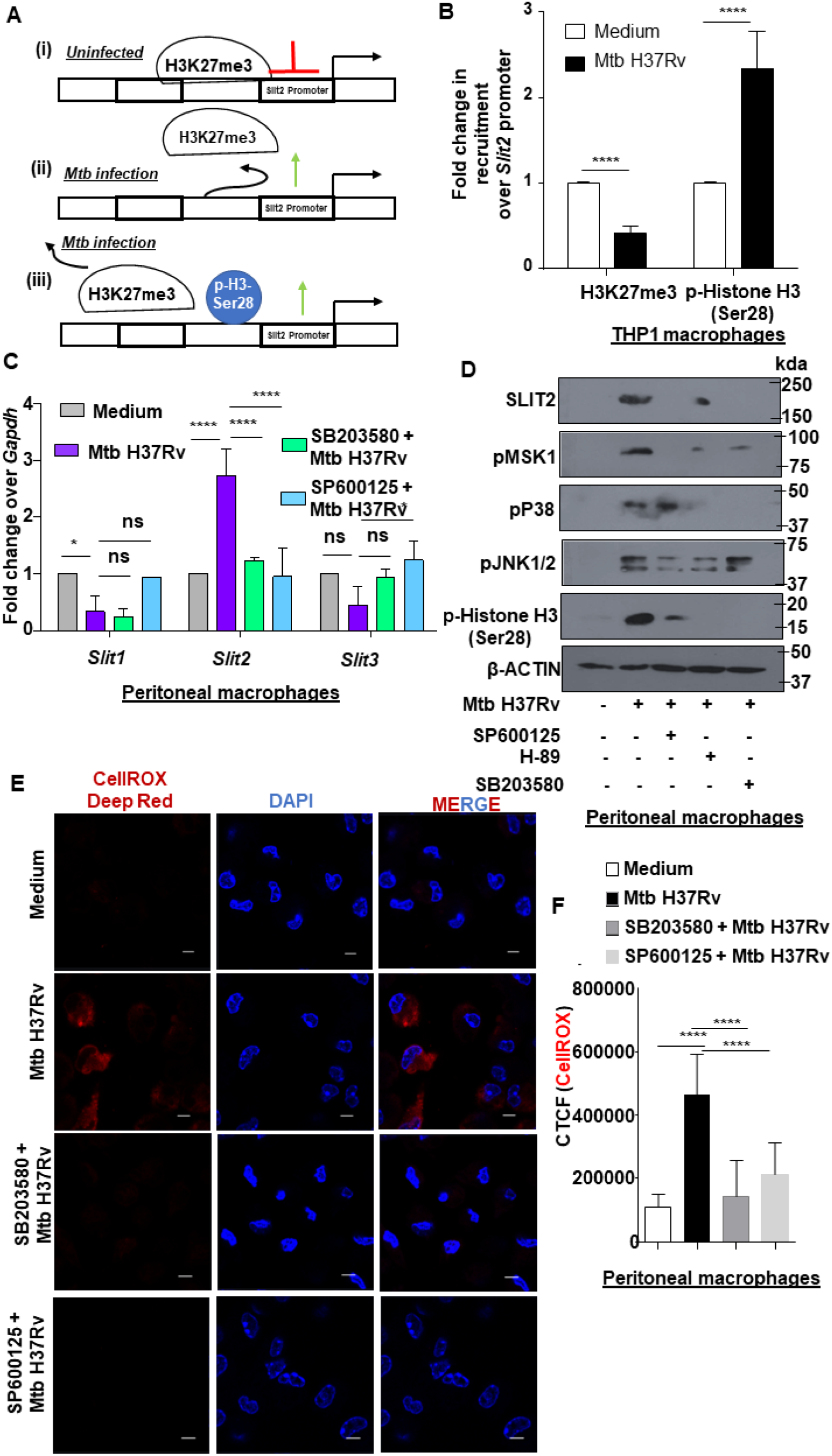
Mtb infection reduces H3K27me3 deposition over Slit2 promoter to augment Slit2 expression. **(A)** Schematic representing the probable mechanism leading to *Slit2* upregulation during Mtb infection. **(B)** THP1 macrophages were infected with Mtb H37Rv for 24 h and assessed for the recruitment of H3K27me3 and p-Histone H3 (Ser28) over the *Slit2* promoter by ChIP assay. **(C)** Mouse peritoneal macrophages were treated with the indicated inhibitors for 1 h, followed by 24 h infection with Mtb H37Rv and assessed for the expression of *Slit* ligands transcripts by qRT-PCR **(D)** Mouse peritoneal macrophages were treated with the indicated inhibitors for 1 h, followed by 24 h infection with Mtb H37Rv and assessed for the expression of indicated proteins by immunoblotting **(E, F)** Mouse peritoneal macrophages were treated with the indicated inhibitors for 1 h. Cells were infected with Mtb H37Rv for 24 h and assessed for the accumulation of ROS by fluorescence microscopy (CellROX Deep Red); **(E)** representative image and **(F)** respective quantification. . All immunoblotting and immunofluorescence data are representative of three independent experiments. β-ACTIN was utilized as loading control. qRT-PCR data represents mean±S.E.M. from three independent experiments. ROS, reactive oxygen species; h, hours; ChIP, Chromatin immunoprecipitation; d, days; kda, kilodalton; CTCF, corrected total cell fluorescence. *, p<0.05; ****, p < 0.0001 (Student’s t-test in B, One-way ANOVA in C,F; GraphPad Prism 6.0 and 9.0). Scale bar, 5 μm. SB203580: p38 inhibitor, SP600125: JNK1/2 inhibitor.

### Vanin1 (VNN1) expression aids SLIT2-dependent ROS accumulation in macrophages

To determine the mechanistic details of ROS accumulation, murine macrophages were treated with recombinant SLIT2 (rSLIT2), and ROS accumulation was assessed using CellROX Deep Red reagent. There was no appreciable increase in the ROS levels **(Fig.S3)**, indicating the possibility of an indirect regulation of ROS by SLIT2 during Mtb infection. Glutathione is a key antioxidant molecule that aids in the maintenance of redox homeostasis(29). Interestingly, the cellular pantetheinase, VNN1 has been reported to regulate antioxidant responses by modulating the levels of glutathione(45, 46). Within a cell, VNN1 forms pantothenic acid and cysteamine, which can be oxidized to the disulfide, cystamine. High concentrations of cystamine inhibit the activity of γ-glutamylcysteine synthetase (γ-GCS) and thereby, contribute to oxidative stress by depleting glutathione(47).

Mtb infection elicited enhanced expression of VNN1, both *in vitro* and *in vivo* **(Fig.3A, Fig.S4A)**. The expression of VNN1 was also amplified in murine macrophages upon rSLIT2 treatment **(Fig.3B, Fig.S4B)**. In line with our observations with SLIT2 expression, perturbation of p38, JNK1/2 or MSK1 phosphorylation compromised VNN1 expression. Besides, inhibition of Protein Kinase A (KT5720), another upstream kinase of MSK1 revealed compromised expression of VNN1 **(Fig.3C)**. Furthermore, Mtb-triggered ROS levels were depleted in cells wherein VNN1 activity was perturbed **(Fig.3D,E; Fig.S4C)**. Glutathione has been reported to augment mycobacterial killing (30, 48). In this direction, we performed an *in vitro* mycobacterial survival assay in murine peritoneal macrophages upon VNN1 inhibition. Interestingly, RR6-mediated VNN1 inhibition minimized mycobacterial survival within macrophages **(Fig.3F)**.

**Figure 3.**
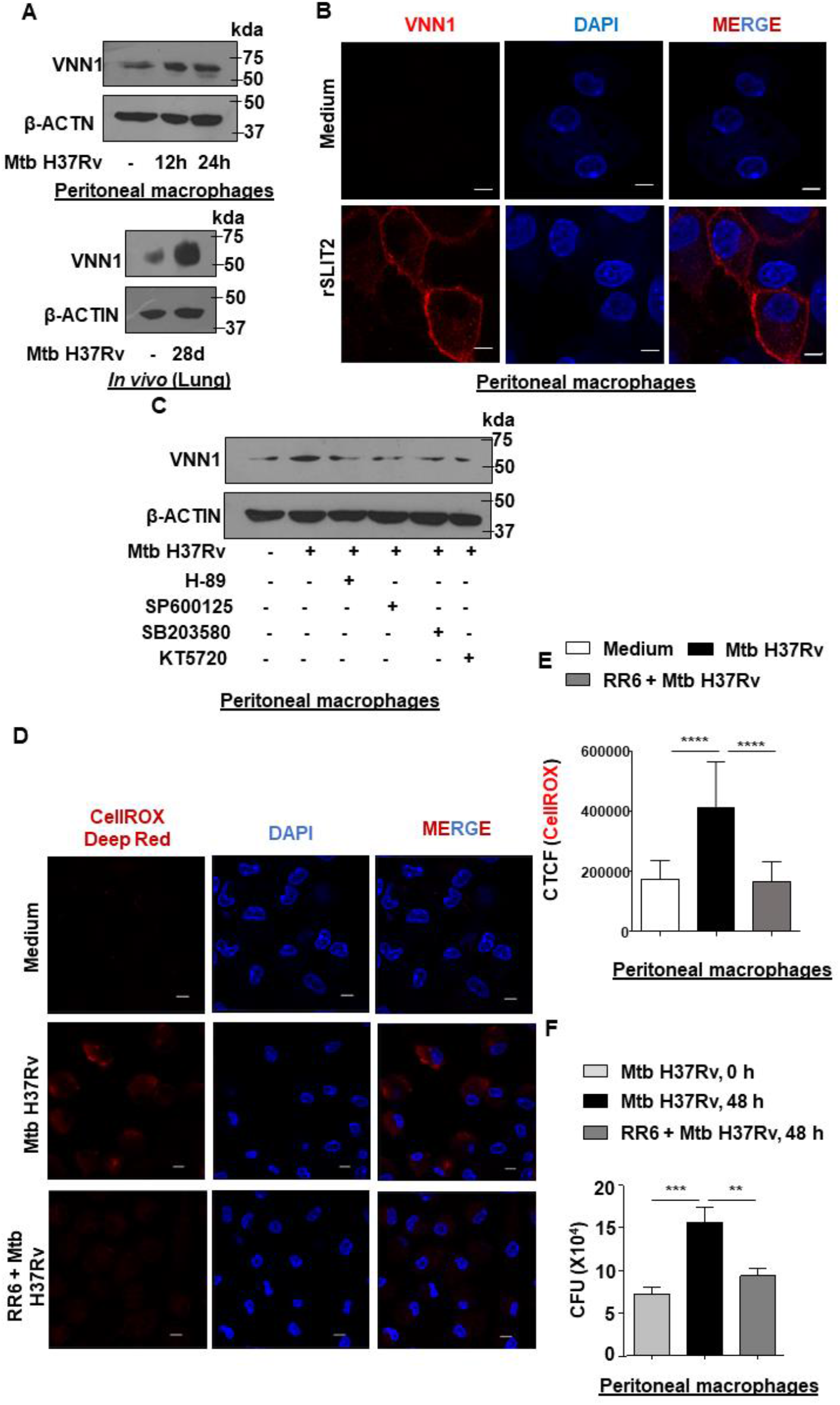
SLIT2-dependent elevated ROS levels is promoted by VNN1 expression. **(A)** Relative expression of VNN1 upon Mtb infection for the indicated time points in mouse peritoneal macrophages *(top)*, and lung homogenates of mice *(below*). (**B)** Mouse peritoneal macrophages were treated with recombinant SLIT2 (rSLIT2) for 12 h and assessed for the expression of VNN1 by immunofluorescence. **(C)** Mouse peritoneal macrophages were treated with the indicated inhibitors for 1 h, followed by 24 h infection with Mtb H37Rv and assessed for the expression of indicated proteins by immunoblotting. **(D,E)** Mouse peritoneal macrophages were treated with VNN1 inhibitor for 1 h. Cells were infected with Mtb H37Rv for 24 h and assessed for the accumulation of ROS by fluorescence microscopy (CellROX Deep Red); **(D)** representative image and **(E)** respective quantification. **(F)** Mouse peritoneal macrophages were infected with Mtb H37Rv for 4 h. Extracellular bacteria were removed, and the infected cells were cultured in the presence or absence of VNN1 inhibitor, RR6 for 48 h. Cells were lysed and plated on 7H11 to enumerate intracellular Mtb H37Rv burden. All immunoblotting and immunofluorescence data are representative of three independent experiments. β-ACTIN was utilized as loading control. ROS, reactive oxygen species; h, hours; d, days; kda, kilodalton; CTCF, corrected total cell fluorescence. **, p<0.01;; *** p<0.001; ****, p < 0.0001 (One way ANOVA in E, F; GraphPad Prism 6.0). Scale bar, 5 μm. RR6: VNN1 inhibitor, SB203580: p38 inhibitor, SP600125: JNK1/2 inhibitor, H-89: MSK1 inhibitor, KT5720: PKA inhibitor

Thus, Mtb infection results in robust expression of VNN1 in a SLIT2-dependent manner which provides survival benefits to the intruding pathogen.

### Aryl hydrocarbon receptor (AhR) contributes to SLIT2-mediated VNN1 expression

Next, we employed a bioinformatics-based approach to scan the promoter region of *Vnn1*, which indicated the putative binding sites for different transcription factors. The presence of Aryl hydrocarbon receptor (AhR) binding sites was intriguing as earlier reports have implicated the AhR pathway in SAX-3/Robo-mediated dorsal migration of neurons in *C. elegans*(49). Initially, we checked the activation of AhR by a luciferase assay-based approach. Specifically, RAW264.7 macrophages were transfected with pGudluc7.5 plasmid which contained AhR binding sites upstream to the promoter of the luciferase gene(50). Transfected cells elicited enhanced luciferase activity upon Mtb infection and rSLIT2 treatment, indicating AhR pathway activation **(Fig.4A)**. Again, reports have implicated AhR in inducing the expression of molecules involved in oxidative stress(51). Particularly, AhR pathway contributed to the increase in superoxide production. In line with these observations, pre-treatment of murine macrophages with AhR inhibitor, CH-223191, compromised VNN1 upregulation upon Mtb infection and rSLIT2 treatment **(Fig.4B)**. Furthermore, CH-223191 pre-treatment to murine macrophages compromised ROS levels upon Mtb infection **(Fig.4C,D)**. Again, perturbation of p38 and JNK1/2 phosphorylation compromised AhR pathway activation as assessed by luciferase assay **(Fig.4E)**. Our results were corroborated further when Mtb-infected macrophages exhibited enhanced recruitment of AhR and its nuclear translocator ARNT on the VNN1 promoter **(Fig.4F)**. Thus, AhR positively regulates Mtb-mediated VNN1 expression.

**Figure 4.**
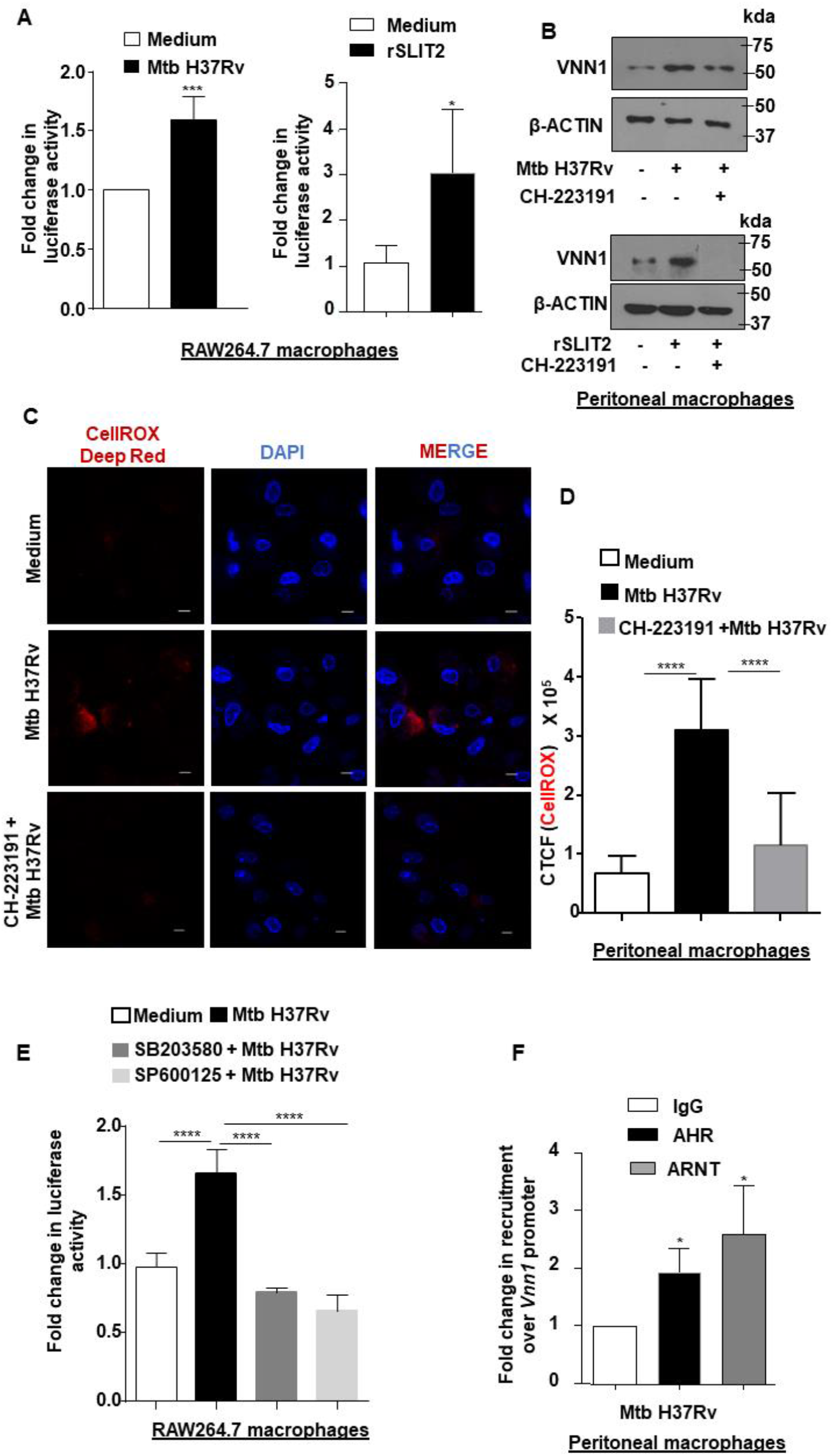
Aryl hydrocarbon receptor contributes to SLIT2-mediated VNN1 expression. **(A)** RAW264.7 macrophages were transiently transfected with pGudLuc7.5 plasmid and the transfected cells were infected with Mtb H37Rv for 24 h *(left side)* or treated with rSLIT2 for 12 h *(right side)*, followed by assessment of luciferase counts using luminometer. **(B)** Mouse peritoneal macrophages were treated with AhR inhibitor for 1 h, followed by 24 h infection with Mtb H37Rv *(top)*, or treated with rSLIT2 for 12 h *(below)* and assessed for the expression of VNN1 by immunoblotting. **(C,D)** Mouse peritoneal macrophages were treated with the AhR inhibitor for 1 h. Cells were infected with Mtb H37Rv for 24 h and assessed for the accumulation of ROS by fluorescence microscopy (CellROX Deep Red); **(C)** representative image and **(D)** respective quantification. **(E)** RAW264.7 macrophages were transiently transfected with pGudLuc7.5 plasmid. Post recovery, the transfected cells were treated with the indicated inhibitors for 1 h, and then infected with Mtb H37Rv for 24 h. Following cell lysis luciferase counts were determined using a luminometer. **(F)** Mouse peritoneal macrophages were infected with Mtb H37Rv for 24 h and assessed for the recruitment of AhR and associated nuclear translocator, ARNT over the *Vnn1* promoter by ChIP assay. All immunoblotting and immunofluorescence data are representative of three independent experiments. β-ACTIN was utilized as loading control. ROS, reactive oxygen species; h, hours; d, days; ChIP, Chromatin immunoprecipitation; kda, kilodalton; CTCF, corrected total cell fluorescence. *, p<0.05; *** p<0.001; ****, p < 0.0001 (Student’s t-test in A,F; One way ANOVA in D,E. GraphPad Prism 6.0 and 9.0). Scale bar, 5 μm. CH-223191: AhR inhibitor

## Conclusion

In the current study, we unravel the interplay of specific signaling components and host epigenetic machinery in governing the pathogenesis of Mtb inside macrophages. While we underscore the role of SLIT2 in Mtb-mediated oxidative stress, it is possible that SLIT2 plays other significant roles in regulating inflammatory responses during TB. It is most pertinent to understand the tissue-specific functions of such molecules. Tissue tolerance to infections define the outcome of disease progression(52). Molecules such as SLIT2 and VNN1 could contribute to enhancing tissue tolerance to the bacteria and thereby favor pathogen persistence(53). Thus, it would be an interesting avenue to explore the potential of SLIT2-a secreted glycoprotein, as a biomarker for intracellular bacterial infections, including TB

Again, we have implicated VNN1 in contributing to enhanced ROS levels within the cells. However, whether enhanced VNN1 activity contributes to depleted glutathione pools by modulating the enzymatic activity of γ -glutamylcysteine synthetase during Mtb infection, needs to be investigated further. This would be challenging as it would involve identification and validation of specific modifications on the said enzyme and correlating them with specific enzyme activity. Besides, the study does not rule out the possibility of modulatory activities of VNN1 on other stress-responsive enzymes such as transglutaminases that would in turn, contribute to imbalance in cellular redox homeostasis(54, 55). Also, the effect of AhR or VNN1 on the expression of other cellular proteins and their potential crosstalk with distinct pathways are important avenues for further research(56, 57). Importantly, VNN1 has been shown to regulate granuloma formation during *Coxiella burnetii* infection(58). We believe that VNN1 may contribute to similar functions during TB pathogenesis. Our *in vitro* results do indicate a beneficial role of VNN1 in mycobacterial survival. Therefore, the regulation of VNN1 may form an effective therapeutic avenue in the context of TB-associated granuloma formation and oxidative functions, and consequently mycobacterial clearance.

Taken together, we report the role of a novel signaling molecule, SLIT2, during Mtb-mediated oxidative stress build-up. Further investigations are necessary to delineate other roles (if any) of this molecule in TB progression and pathology.

## Materials and Methods

### Mice

BALB/c mice (male or female/ 4-6 weeks) were utilized for all experiments. Mice were purchased from The Jacksons Laboratory and maintained at the Central Animal Facility (CAF) in the Indian Institute of Science (IISc) under 12-h light and dark cycle.

### Ethics Statement

Requisite approval from Institutional Ethics Committee for animal experimentation was obtained. The animal care and use protocol adhered were approved by national guidelines of the Committee for the Purpose of Control and Supervision of Experiments on Animals (CPCSEA), Government of India. Experiments with virulent mycobacteria (Mtb H37Rv) were approved by the Institutional Biosafety Committee.

### Peritoneal macrophages

For *in vitro* experiments, mouse peritoneal macrophages were utilized. Briefly, mice were injected intraperitoneally with (4-8 %) Brewer’s thioglycollate and peritoneal exudates were harvested in ice cold PBS after four days, seeded in tissue culture dishes. Cells were cultured in DMEM with 10% FBS supplement for 8-12 h. Adherent cells were utilized as peritoneal macrophages.

### Bone marrow-derived macrophages

Tibia and femur from BALB/c mice were flushed with ice-cold DMEM containing 10% fetal bovine serum. Bone marrow was collected in 50ml tube and bone marrow clusters were disintegrated by pipetting. The cell suspension was centrifuged (1500rpm) for 5min at 4°C. Thereafter, the cells were washed with DMEM containing 10% fetal bovine serum. RBC was lysed by treating the cells with RBC lysis buffer followed by two washes with DMEM containing 10% fetal bovine serum. The cells were seeded in DMEM containing 10% fetal bovine serum and 20% of LN29 cell supernatant. The cells were incubated at 37°C, 5% CO_2_ and 95% humidity in a CO _2_ incubator. Medium supplementation was done on the third and fifth day with DMEM containing 10% fetal bovine serum and 20% LN29 cell supernatant. After seven days of differentiation, the cells were used for further experiments.

### Cells

RAW 264.7 mouse monocyte-like cell line and human monocytic cell line THP1 cells were obtained from American Type Culture Collection (ATCC), USA. RAW264.7 cells were cultured in Dulbecco’s Modified Eagle Medium (DMEM, Gibco, Thermo Fisher Scientific) supplemented with 10% heat inactivated Fetal Bovine Serum (FBS, Gibco, Thermo Fisher Scientific) and maintained at 37 °C in 5% CO_2_ incubator. THP1 cells were cultured in RPMI-1640 synthetic medium with 10% fetal bovine serum (FBS) supplement. Monocytes were differentiated by treatment with 20 ng/mL phorbol 12-myristate 13-acetate (PMA) for 18 h. Thereafter, cells were rested for two days to ensure their reversion to a resting phenotype before infection.

### Bacteria

Virulent strain of *Mycobacterium tuberculosis* (Mtb H37Rv) was a kind research gift from Prof. Amit Singh, Department of Microbiology and Cell Biology, and Centre for Infectious Disease Research, IISc. Briefly, single colonies of mycobacteria were cultured in Middlebrook 7H9 medium (Difco, USA) supplemented with 10% OADC (oleic acid, albumin, dextrose, catalase). Single-cell suspensions of the bacteria were obtained by passing mid-log phase culture through 23-, 28- and 30-gauge needle, ten times each. These suspensions were used for infecting primary cells or RAW 264.7 cells at multiplicity of infection 10. The studies involving virulent mycobacterial strains were carried out in the biosafety level 3 (BSL-3) facility at the Centre for Infectious Disease Research (CIDR), IISc.

### Reagents

All general chemicals and reagents were procured from Sigma-Aldrich/ Merck Millipore, and Promega. Tissue culture plastic ware was purchased from Jet Biofil and Corning Inc. siRNAs were obtained from Dharmacon as siGENOME SMART-pool reagents against *Slit2*, *Vnn1*. Oleic acid, 4′,6-Diamidino-2-phenylindole dihydrochloride (DAPI) were procured from Sigma-Aldrich. Lipofectamine 3000 was purchased from Thermo Fisher Scientific.

### Antibodies

| Antibody | Company | Catalog No. |
| --- | --- | --- |
| SLIT2 | GeneTex | GTX118220 |
| ROBO2 | GeneTex | GTX55278 |
| p-Histone H3 (Ser 28) | CST | 9713 |
| p-MSK1 (Ser376) | CST | 9591 |
| p-P38 (Thr180/Tyr182) | CST | 9211 |
| p-JNK 1/2 (Thr183/Tyr185) | CST | 4671 |
| VNN1 | Santa Cruz | sc-23907 |
| VNN1 | Abcam | ab205912 |
| AHR | Abcam | ab84833 |
| ARNT | CST | 5537 |
| HRP-tagged anti-β-ACTIN | Sigma | A3854 |
| HRP conjugated anti-rabbit IgG | Jackson ImmunoResearch (USA) | 111-035-045 |

### Treatment with pharmacological reagents

Primary macrophages pre-treated with the following reagents one hour prior to infection with Mtb H37Rv: SB203580 (Calbiochem, 10 μM); SP600125 (Calbiochem, 10 μM); H89 (Calbiochem, 10 μM); KT5720 (Calbiochem, 10 μM); RR6 (MedChemExpress 20 μM); CH-223191(Calbiochem, 10 μM). Recombinant SLIT2 (rSLIT2) was added as indicated (R&D Systems, 100ng/mL).

### Plasmids

β-galactosidase plasmid was received as kind gift from Prof. Kumaralvel Somasundaram, (IISc, Bangalore). pGudLuc7.5 was a kind gift from a kind gift from Prof. Denison MS, University of California, Davis, California.

### Transient transfection studies

RAW 264.7 macrophages were transfected with the indicated plasmids; or primary macrophages were transfected with 100 nM each of siGLO Lamin A/C, non-targeting siRNA or specific siRNAs with the help of Lipofectamine 3000 for 6 h; followed by 24 h recovery. 70-80 % transfection efficiency was observed by counting the number of siGLO Lamin A/C positive cells in a microscopic field using fluorescence microscopy. Transfected cells were subjected to the required infections/ treatments for the indicated time points and processed for analyses.

### Luciferase Assay

RAW 264.7 cells were transfected with pGudLuc7.5 and β-galactosidase plasmids using Lipofectamine 3000 for 6 h, followed by 24 h of recovery. The transfected cells were subjected to the required infections/ treatments for the indicated time points and processed for analyses. Briefly, cells were harvested and lysed in reporter lysis buffer (Promega) and luciferase activity was assayed using luciferase assay reagent (Promega). The results were normalized for transfection efficiencies by assay of β-galactosidase activity

### RNA isolation and quantitative real time PCR (qRT-PCR)

Treated samples were harvested in TRl-Reagent (Sigma-Aldrich) and incubated with chloroform for phase separation. Total RNA was precipitated from the aqueous layer. Equal amount of RNA was converted into cDNA using First Strand cDNA synthesis kit (Applied Biological Materials Inc.). The obtained cDNA was used for SYBR Green (Thermo Fisher Scientific) based quantitative real time PCR analysis for the concerned genes. *Gapdh* was used as internal control gene. Primer pairs used for expression analyses are provided below:

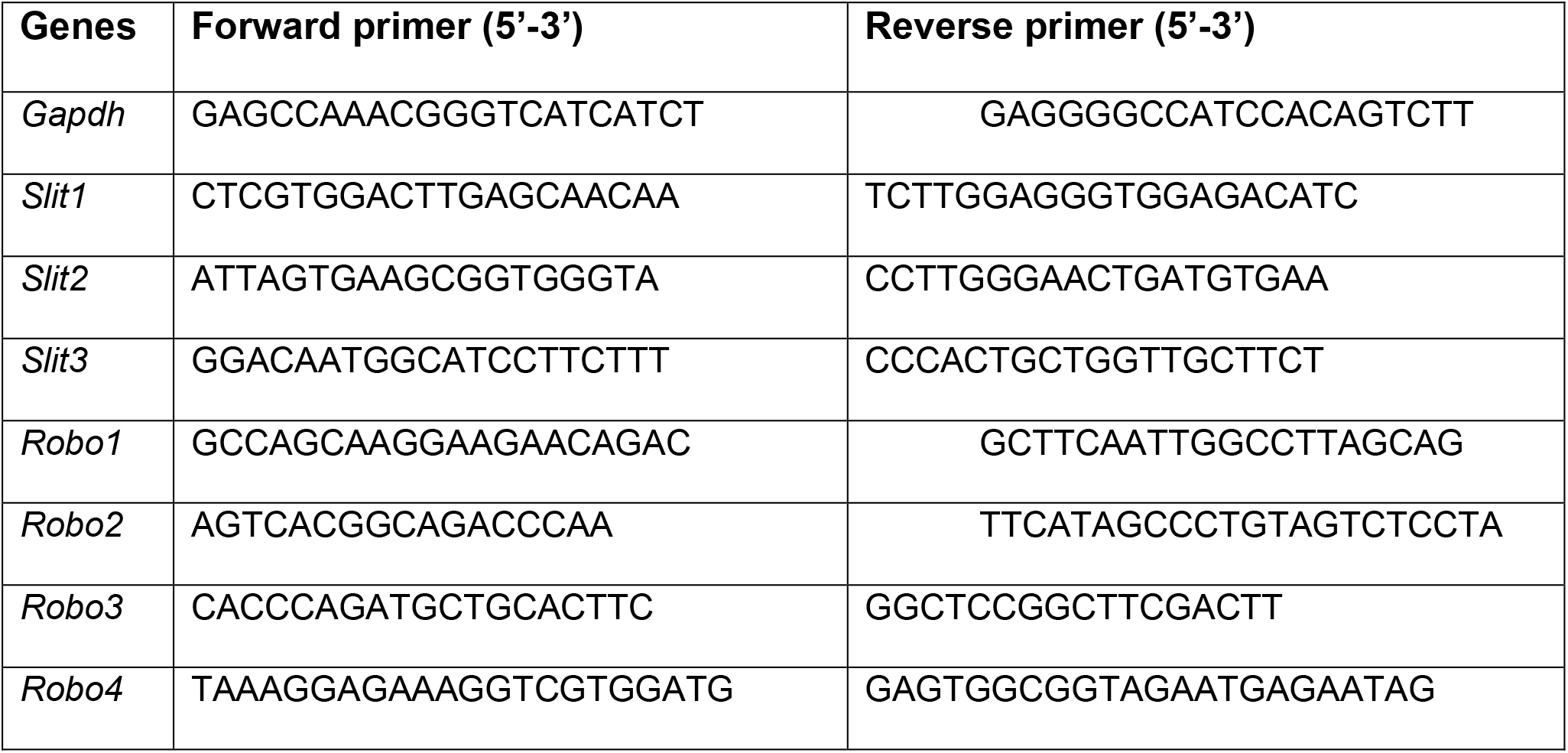

Primers were synthesized and obtained from Eurofins Genomics Pvt. Ltd. (India).

### Immunoblotting

For immunoblotting, cells were washed with PBS after treatment/ infection. Whole cell lysate was prepared by lysing in RIPA buffer [50 mM Tris-HCl (pH 7.4), 1% NP-40, 0.25% sodium deoxycholate, 150 mM NaCl, 1 mM EDTA, 1 mM PMSF, 1 μg/ml each of aprotinin, leupeptin, pepstatin, 1 mM Na_3_VO_4_, 1 mM NaF] on ice for 30 min. The total protein from cell lysates was estimated by Bradford reagent. Subsequently, equal amount of protein was resolved on 12% or 10% SDS-PAGE and transferred onto PVDF membranes (Millipore) by semi-dry immunoblotting method (Bio-Rad). 5% non-fat dry milk powder in TBST [20 mM Tris-HCl (pH 7.4), 137 mM NaCl, and 0.1% Tween 20] was used for blocking nonspecific binding for 60 min at room temperature. After washing with TBST, the blots were incubated overnight at 4°C with primary antibody diluted in TBST with 5% BSA. After washing with TBST, blots were incubated with anti-rabbit IgG secondary antibody conjugated to HRP antibody for 4h at 4°C. The immunoblots were developed with enhanced chemiluminescence detection system (Perkin Elmer) as per manufacturer’s instructions. For developing more than one protein at a particular molecular weight range, the blots were stripped off the first antibody at 60 °C for 5 min using stripping buffer (62.5 mM Tris-HCl, with 2 % SDS, 100 mM 2-Mercaptoethanol), washed with 1X TBST, blocked (as mentioned above); followed by probing with the subsequent antibody following the described procedure. β-ACTIN was used as loading control.

### Chromatin Immunoprecipitation (ChIP) Assay

The experiments involving ChIP assays were carried out using a protocol provided by Upstate Biotechnology, Sigma-Aldrich and Abcam with certain modifications. Briefly, treated samples were washed with ice cold PBS and fixed with 3.6 % formaldehyde for 15 min. at room temperature followed by inactivation of formaldehyde with 125 mM glycine. Nuclei were lysed in 0.1% SDS lysis buffer [50 mM Tris-HCl (pH 8.0), 200 mM NaCl, 10 mM HEPES (pH 6.5), 0.1 % SDS, 10 mM EDTA, 0.5 mM EGTA, 1 mM PMSF, 1 μg/ml of each aprotinin, leupeptin, pepstatin, 1 mM Na_3_VO_4_ and 1 mM NaF]. Chromatin was sheared in the Bioruptor Plus (Diagenode, Belgium) at high power for 60 rounds of 30 sec pulse ON and 45 sec pulse OFF. Chromatin extracts containing DNA fragments with an average size of 500 bp were immunoprecipitated with the indicated antibodies or rabbit preimmune sera complexed with Protein A agarose beads (Sigma-Aldrich). Immunoprecipitated complexes were sequentially washed with Wash Buffer A, B and TE [Wash Buffer A: 50 mM Tris-HCl (pH 8.0), 500 mM NaCl, 1 mM EDTA, 1 % Triton X-100, 0.1 % Sodium deoxycholate, 0.1 % SDS and protease/phosphatase inhibitors; Wash Buffer B: 50 mM Tris-HCl (pH 8.0), 1 mM EDTA, 250 mM LiCl, 0.5 % NP-40, 0.5 % Sodium deoxycholate and protease/phosphatase inhibitors; TE: 10 mM Tris-HCl (pH 8.0), 1 mM EDTA] and eluted in elution buffer [1 % SDS, 0.1 M NaHCO_3_]. After treating the eluted samples with RNase A and Proteinase K, DNA was purified and precipitated using phenol-chloroform-ethanol method. Purified DNA was analyzed by quantitative real time RT-PCR. All values in the test samples were normalized to amplification of the specific gene in Input and IgG pull down and represented as fold change in modification or enrichment. All ChIP experiments were repeated at least three times. The list of primers is given below:

Primers were synthesized and obtained from Eurofins Genomics Pvt. Ltd. (India).

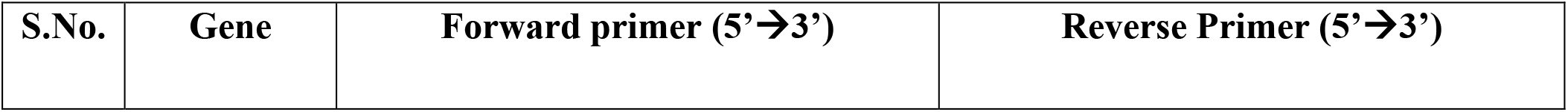
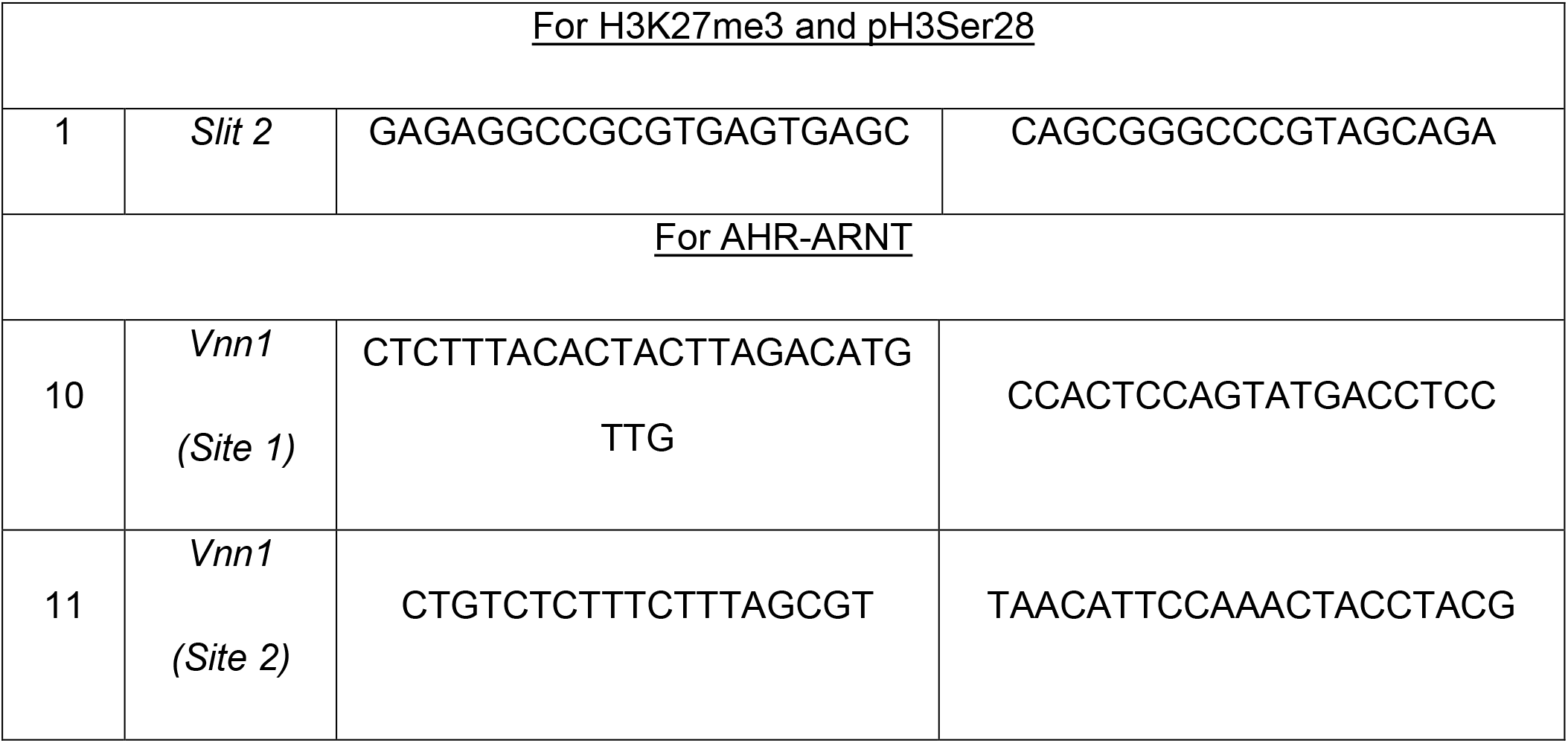

### *In vivo* infection of mouse

BALB/c mice (n=5) were infected with mid-log phase Mtb H37Rv, using a Madison chamber aerosol generation instrument calibrated to 100 CFU/animal. Aerosolized animals were maintained in securely commissioned BSL3 facility. Post 28 days of established infection, mice were sacrificed. Specific lobes from the lungs of mice were homogenized for the extraction of RNA and protein

### CellROX staining for determining oxidative stress

CellROX Deep Red Reagent (Thermo Fisher Scientific, USA) was used to determine oxidative stress in macrophages as per manufacturer’s instructions. Briefly, macrophages were treated with CellROX Deep Red Reagent at a final concentration of 5 μM, diluted in DMEM without phenol red. The treated cells were incubated for 30 min at 37 °C in the CO_2_ incubator. Cells were then washed with PBS thrice, followed by fixation with 3.6 % formaldehyde for 15 min. Nuclei were stained with DAPI and images were captured in Zeiss LSM 710 confocal laser scanning microscope.

### *In vitro* CFU analysis

Mouse peritoneal macrophages were infected with Mtb H37Rv at MOI 5 for 4 h. Thereafter, the cells were thoroughly washed with PBS to remove any surface adhered bacteria. The cells were incubated in medium containing amikacin (0.2 mg/mL) 2 h to deplete any extracellular mycobacteria. After amikacin treatment, the cells were thoroughly washed with PBS and taken for 0 h time point and a duplicate set was maintained in antibiotic free medium for next 48 h along with VNN1 inhibitor, as indicated. Intracellular mycobacterial burden was enumerated by lysing macrophages with 0.06 % SDS in 7H9 Middlebrook medium. Appropriate dilutions were plated on Middlebrook 7H11 agar plates supplemented with OADC (oleic acid, albumin, dextrose, catalase). Total colony forming units (CFUs) were counted after 21 days of plating.

#### Statistical analysis

Levels of significance for comparison between samples were determined by the student’s t-test and one-way ANOVA followed by Tukey’s multiple-comparisons. The data in the graphs are expressed as the mean ± S.E. for the values from at least 3 or more independent experiments and P values < 0.05 were defined as significant. GraphPad Prism software (6.0 and 9.0 versions, GraphPad Software, USA) was used for all the statistical analyses.

## Acknowledgements

We thank CAF, IISc for maintaining and providing mice for experimentation. β-galactosidase plasmid was a kind research gift from Prof. Kumaravel Somasundaram, Department of Microbiology and Cell Biology, IISc. pGudLuc7.5 plasmid was a kind gift from Prof. Denison MS, University of California, Davis, California. We acknowledge the help of BSL-3 facility and staff for helping us in our *in vitro* and *in vivo* experiments with Mtb H37Rv. We are grateful to Prof. Balaji K.N.’s research group, majorly Mahima, Ankita Ghoshal, Awantika Shah, and Smriti Sundar for their valuable inputs in improving the manuscript. We thank Dr. Tanushree Mukherjee for her critical comments on the manuscript and help with data analysis.

## Author contributions

SMB, conceptualization, investigation, formal analysis, manuscript draft preparation, editing; SB, investigation, formal analysis; KNB, conceptualization, formal analysis, supervision, manuscript review and editing.

---

**Figure S1.**
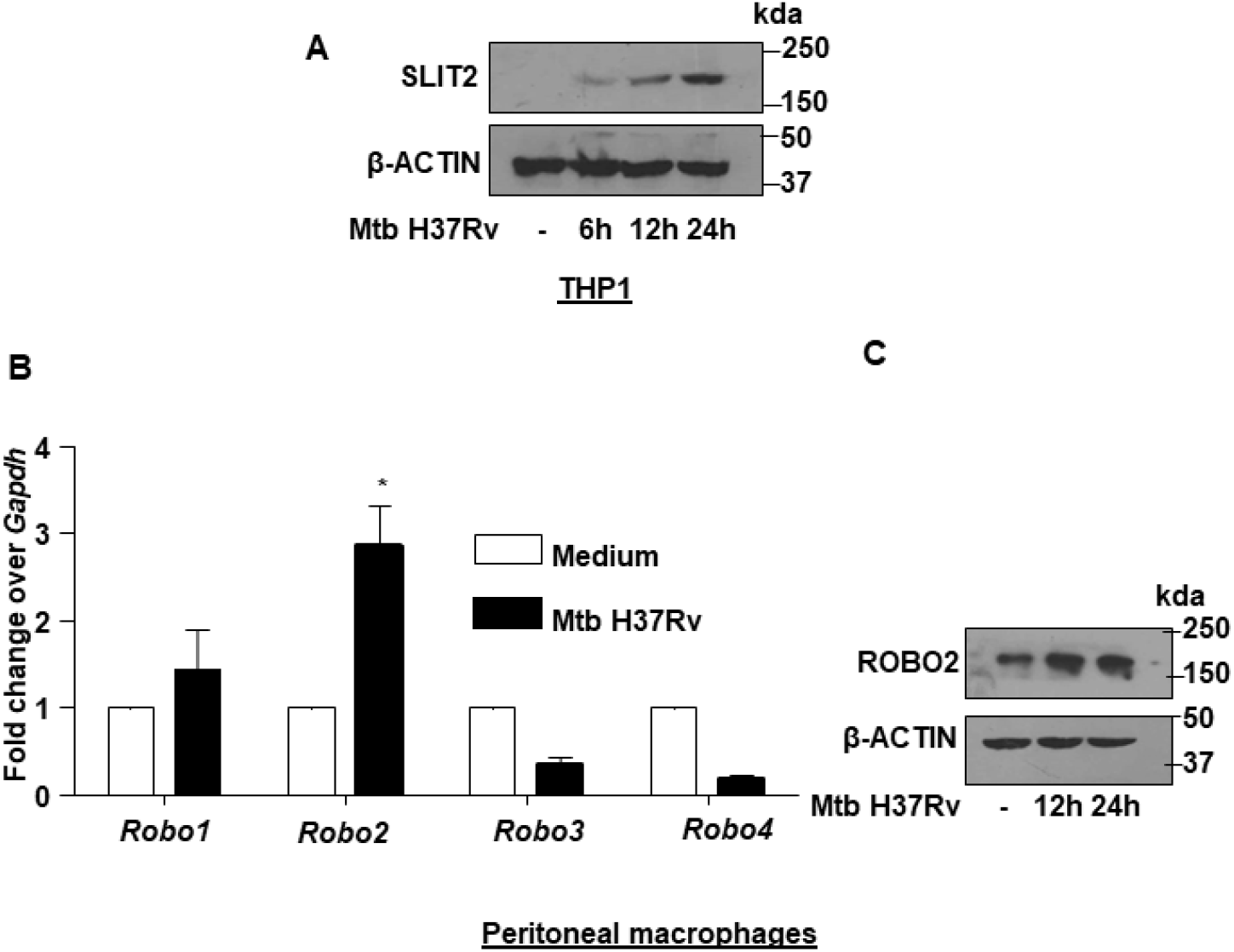
**(A)** THP1 macrophages were infected with Mtb H37Rv for the indicated time points. Whole cell lysates were assessed for SLIT2 protein expression by immunoblotting. **(B)** Mouse peritoneal macrophages were infected with Mtb H37Rv for 24 h and assessed for the transcript levels of ROBO receptors. **(C)** Mouse peritoneal macrophages were infected with Mtb H37Rv for the indicated time points and ROBO2 levels were assessed by immunoblotting. All immunoblotting data are representative of three independent experiments. β-ACTIN was utilized as loading control. qRT-PCR data represents mean±S.E.M. from three independent experiments.

**Figure S2.**
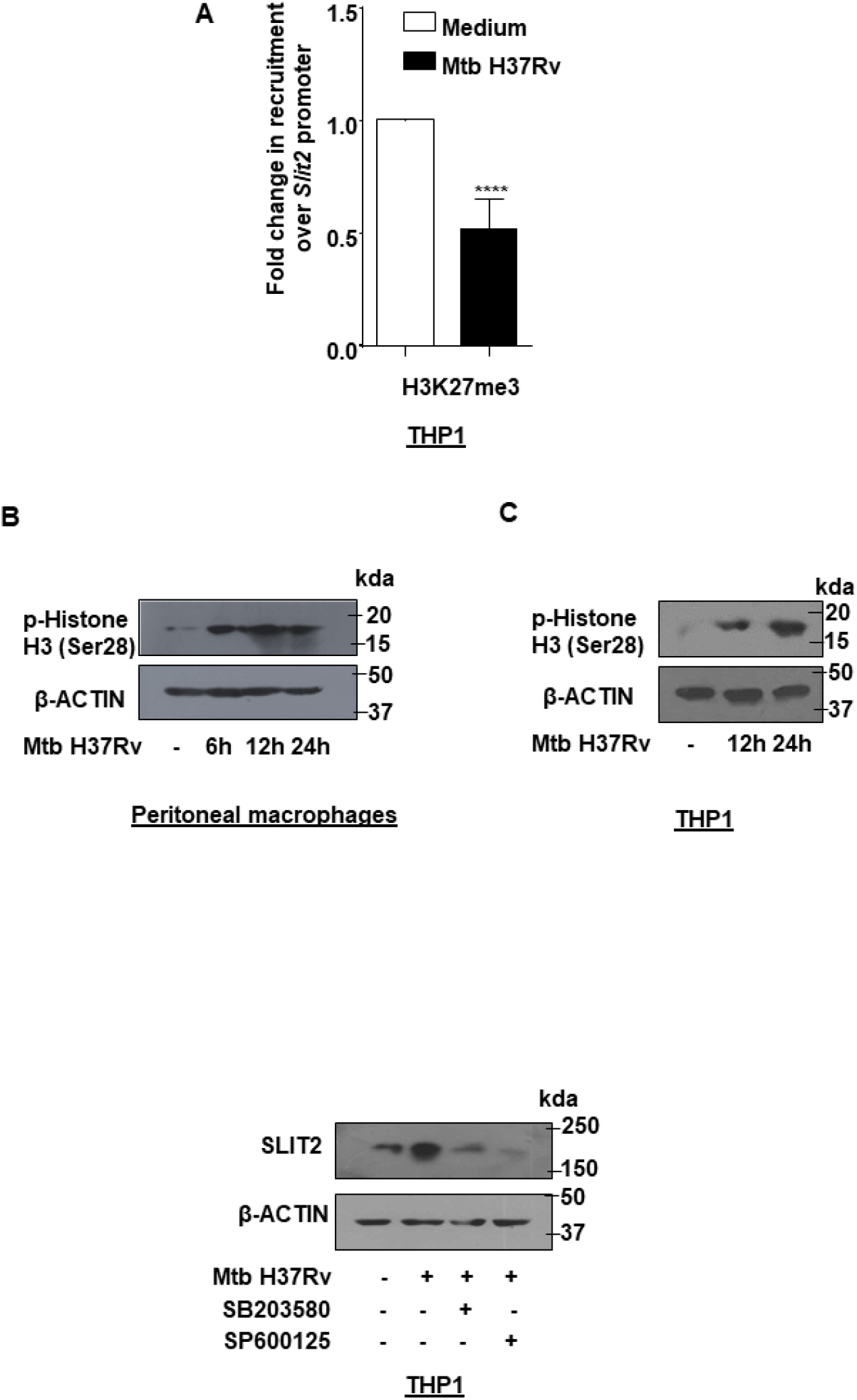
**(A)** THP1 macrophages were infected with Mtb H37Rv for 24 h and assessed for the recruitment of H3K27me3 over the *Slit2* promoter by ChIP assay. **(B)** Mouse peritoneal macrophages were infected with Mtb H37Rv for the indicated time points and assessed for the levels of p-Histone H3 (Ser28) by immunoblotting. **(C)** THP1 macrophages were infected with Mtb H37Rv for the indicated time points and assessed for the levels of p-Histone H3 (Ser28) by immunoblotting. **(D)** THP1 macrophages were treated with the indicated inhibitors for 1 h, followed by 24 h infection with Mtb H37Rv and assessed for the expression of SLIT2 by immunoblotting. All immunoblotting data are representative of three independent experiments. β-ACTIN was utilized as loading control.

**Figure S3.**
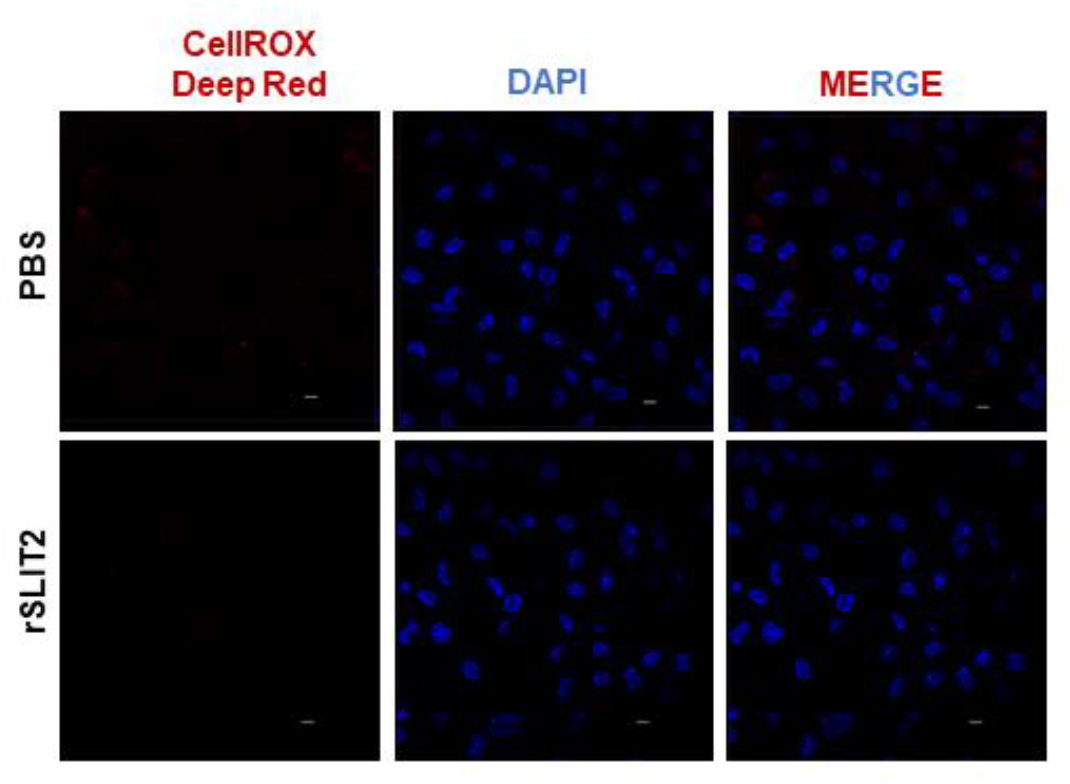
Mouse peritoneal macrophages were treated with rSLIT2 for 12 h and assessed for the accumulation of ROS by fluorescence microscopy (CellROX Deep Red). Scale bar, 5 μm.

**Figure S4.**
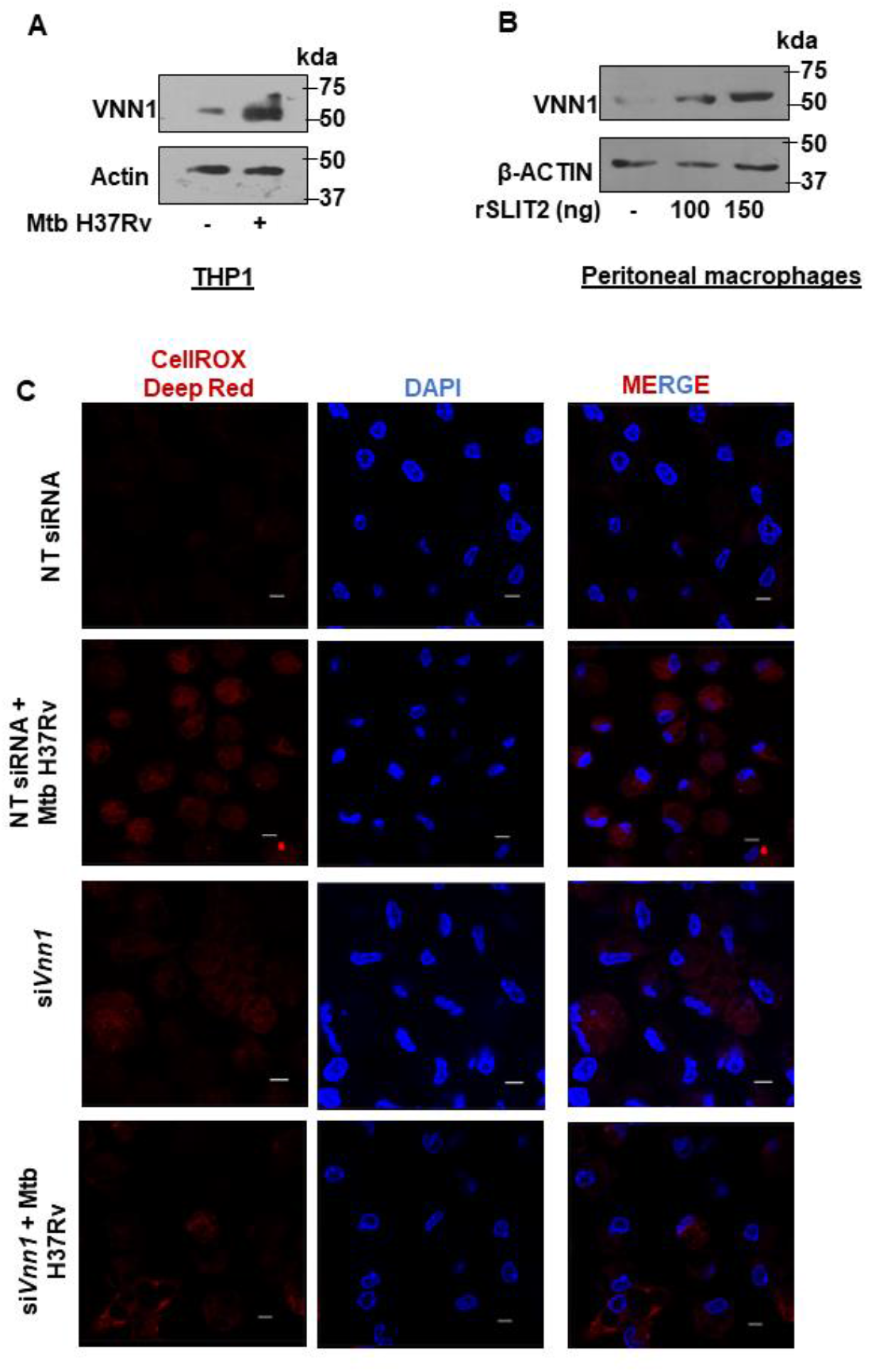
**(A)** THP1 macrophages were infected with Mtb H37Rv for 24 h and assessed for the levels of VNN1 by immunoblotting. **(B)** Mouse peritoneal macrophages were treated with rSLIT2 at the indicated concentrations for 12 h and assessed for the levels of VNN1 by immunoblotting. **(C)** Mouse peritoneal macrophages were transfected with NT or *Vnn1* siRNAs. Transfected cells were infected with Mtb H37Rv for 24 h and assessed for the accumulation of ROS by fluorescence microscopy (CellROX Deep Red). All immunoblotting and immunofluorescence data are representative of three independent experiments. β-ACTIN was utilized as loading control. Scale bar, 5 μm.

## References

1. Ravesloot-Chavez MM, Van Dis E, Stanley SA. The Innate Immune Response to Mycobacterium tuberculosis Infection. Annu Rev Immunol. 2021;39:611–37.

2. Stutz MD, Clark MP, Doerflinger M, Pellegrini M. Mycobacterium tuberculosis: Rewiring host cell signaling to promote infection. J Leukoc Biol. 2018;103(2):259–68.

3. Holla S, Prakhar P, Singh V, Karnam A, Mukherjee T, Mahadik K, et al. MUSASHI-Mediated Expression of JMJD3, a H3K27me3 Demethylase, Is Involved in Foamy Macrophage Generation during Mycobacterial Infection. PLoS Pathog. 2016;12(8):e1005814.

4. Kapoor N, Narayana Y, Patil SA, Balaji KN. Nitric oxide is involved in Mycobacterium bovis bacillus Calmette-Guerin-activated Jagged1 and Notch1 signaling. J Immunol. 2010;184(6):3117–26.

5. Mukherjee T, Balaji KN. The WNT Framework in Shaping Immune Cell Responses During Bacterial Infections. Front Immunol. 2019;10:1985.

6. Holla S, Kurowska-Stolarska M, Bayry J, Balaji KN. Selective inhibition of IFNG-induced autophagy by Mir155- and Mir31-responsive WNT5A and SHH signaling. Autophagy. 2014;10(2):311–30.

7. Liu F, Chen J, Wang P, Li H, Zhou Y, Liu H, et al. MicroRNA-27a controls the intracellular survival of Mycobacterium tuberculosis by regulating calcium-associated autophagy. Nat Commun. 2018;9(1):4295.

8. Leisching GR. Susceptibility to Tuberculosis Is Associated With PI3K-Dependent Increased Mobilization of Neutrophils. Front Immunol. 2018;9:1669.

9. Manca C, Tsenova L, Freeman S, Barczak AK, Tovey M, Murray PJ, et al. Hypervirulent M. tuberculosis W/Beijing strains upregulate type I IFNs and increase expression of negative regulators of the Jak-Stat pathway. J Interferon Cytokine Res. 2005;25(11):694–701.

10. Ting LM, Kim AC, Cattamanchi A, Ernst JD. Mycobacterium tuberculosis inhibits IFN-gamma transcriptional responses without inhibiting activation of STAT1. J Immunol. 1999;163(7):3898–906.

11. Blockus H, Chedotal A. Slit-Robo signaling. Development. 2016;143(17):3037–44.

12. Ypsilanti AR, Zagar Y, Chedotal A. Moving away from the midline: new developments for Slit and Robo. Development. 2010;137(12):1939–52.

13. Wu MF, Liao CY, Wang LY, Chang JT. The role of Slit-Robo signaling in the regulation of tissue barriers. Tissue Barriers. 2017;5(2):e1331155.

14. Li GJ, Yang Y, Yang GK, Wan J, Cui DL, Ma ZH, et al. Slit2 suppresses endothelial cell proliferation and migration by inhibiting the VEGF-Notch signaling pathway. Mol Med Rep. 2017;15(4):1981–8.

15. Wang B, Xiao Y, Ding BB, Zhang N, Yuan X, Gui L, et al. Induction of tumor angiogenesis by Slit-Robo signaling and inhibition of cancer growth by blocking Robo activity. Cancer Cell. 2003;4(1):19–29.

16. Wu JY, Feng L, Park HT, Havlioglu N, Wen L, Tang H, et al. The neuronal repellent Slit inhibits leukocyte chemotaxis induced by chemotactic factors. Nature. 2001;410(6831):948–52.

17. Zhou X, Yao Q, Sun X, Gong X, Yang Y, Chen C, et al. Slit2 ameliorates renal inflammation and fibrosis after hypoxia-and lipopolysaccharide-induced epithelial cells injury in vitro. Exp Cell Res. 2017;352(1):123–9.

18. Terzi A, Roeder H, Weaver CJ, Suter DM. Neuronal NADPH oxidase 2 regulates growth cone guidance downstream of slit2/robo2. Dev Neurobiol. 2021;81(1):3–21.

19. Zhao H, Anand AR, Ganju RK. Slit2-Robo4 pathway modulates lipopolysaccharide-induced endothelial inflammation and its expression is dysregulated during endotoxemia. J Immunol. 2014;192(1):385–93.

20. Pajuelo D, Gonzalez-Juarbe N, Niederweis M. NAD hydrolysis by the tuberculosis necrotizing toxin induces lethal oxidative stress in macrophages. Cell Microbiol. 2020;22(1):e13115.

21. Ravimohan S, Kornfeld H, Weissman D, Bisson GP. Tuberculosis and lung damage: from epidemiology to pathophysiology. Eur Respir Rev. 2018;27(147).

22. Amaral EP, Vinhaes CL, Oliveira-de-Souza D, Nogueira B, Akrami KM, Andrade BB. The Interplay Between Systemic Inflammation, Oxidative Stress, and Tissue Remodeling in Tuberculosis. Antioxid Redox Signal. 2021;34(6):471–85.

23. Braunstein M, Espinosa BJ, Chan J, Belisle JT, Jacobs WR, Jr. SecA2 functions in the secretion of superoxide dismutase A and in the virulence of Mycobacterium tuberculosis. Mol Microbiol. 2003;48(2):453–64.

24. Kurtz S, McKinnon KP, Runge MS, Ting JP, Braunstein M. The SecA2 secretion factor of Mycobacterium tuberculosis promotes growth in macrophages and inhibits the host immune response. Infect Immun. 2006;74(12):6855–64.

25. Shastri MD, Shukla SD, Chong WC, Dua K, Peterson GM, Patel RP, et al. Role of Oxidative Stress in the Pathology and Management of Human Tuberculosis. Oxid Med Cell Longev. 2018;2018:7695364.

26. Virag L, Jaen RI, Regdon Z, Bosca L, Prieto P. Self-defense of macrophages against oxidative injury: Fighting for their own survival. Redox Biol. 2019;26:101261.

27. Costa DL, Namasivayam S, Amaral EP, Arora K, Chao A, Mittereder LR, et al. Pharmacological Inhibition of Host Heme Oxygenase-1 Suppresses Mycobacterium tuberculosis Infection In Vivo by a Mechanism Dependent on T Lymphocytes. mBio. 2016;7(5).

28. Lin K, O'Brien KM, Trujillo C, Wang R, Wallach JB, Schnappinger D, et al. Mycobacterium tuberculosis Thioredoxin Reductase Is Essential for Thiol Redox Homeostasis but Plays a Minor Role in Antioxidant Defense. PLoS Pathog. 2016;12(6):e1005675.

29. Aquilano K, Baldelli S, Ciriolo MR. Glutathione: new roles in redox signaling for an old antioxidant. Front Pharmacol. 2014;5:196.

30. Venketaraman V, Millman A, Salman M, Swaminathan S, Goetz M, Lardizabal A, et al. Glutathione levels and immune responses in tuberculosis patients. Microb Pathog. 2008;44(3):255–61.

31. Russell SA, Bashaw GJ. Axon guidance pathways and the control of gene expression. Dev Dyn. 2018;247(4):571–80.

32. Oswald MCW, Garnham N, Sweeney ST, Landgraf M. Regulation of neuronal development and function by ROS. FEBS Lett. 2018;592(5):679–91.

33. Pocock R, Hobert O. Oxygen levels affect axon guidance and neuronal migration in Caenorhabditis elegans. Nat Neurosci. 2008;11(8):894–900.

34. Dallol A, Krex D, Hesson L, Eng C, Maher ER, Latif F. Frequent epigenetic inactivation of the SLIT2 gene in gliomas. Oncogene. 2003;22(29):4611–6.

35. Dunwell TL, Dickinson RE, Stankovic T, Dallol A, Weston V, Austen B, et al. Frequent epigenetic inactivation of the SLIT2 gene in chronic and acute lymphocytic leukemia. Epigenetics. 2009;4(4):265–9.

36. Jin J, You H, Yu B, Deng Y, Tang N, Yao G, et al. Epigenetic inactivation of SLIT2 in human hepatocellular carcinomas. Biochem Biophys Res Commun. 2009;379(1):86–91.

37. Zhang TJ, Xu ZJ, Wen XM, Gu Y, Ma JC, Yuan Q, et al. SLIT2 promoter hypermethylation-mediated SLIT2-IT1/miR-218 repression drives leukemogenesis and predicts adverse prognosis in myelodysplastic neoplasm. Leukemia. 2022;36(10):2488–98.

38. Yu J, Cao Q, Yu J, Wu L, Dallol A, Li J, et al. The neuronal repellent SLIT2 is a target for repression by EZH2 in prostate cancer. Oncogene. 2010;29(39):5370–80.

39. Sarmento OF, Svingen PA, Xiong Y, Sun Z, Bamidele AO, Mathison AJ, et al. The Role of the Histone Methyltransferase Enhancer of Zeste Homolog 2 (EZH2) in the Pathobiological Mechanisms Underlying Inflammatory Bowel Disease (IBD). J Biol Chem. 2017;292(2):706–22.

40. Xu WD, Huang Q, Huang AF. Emerging role of EZH2 in rheumatic diseases: A comprehensive review. Int J Rheum Dis. 2022.

41. Baek SH. When signaling kinases meet histones and histone modifiers in the nucleus. Mol Cell. 2011;42(3):274–84.

42. Rossetto D, Avvakumov N, Cote J. Histone phosphorylation: a chromatin modification involved in diverse nuclear events. Epigenetics. 2012;7(10):1098–108.

43. Roux PP, Blenis J. ERK and p38 MAPK-activated protein kinases: a family of protein kinases with diverse biological functions. Microbiol Mol Biol Rev. 2004;68(2):320–44.

44. Iida C, Ohsawa S, Taniguchi K, Yamamoto M, Morata G, Igaki T. JNK-mediated Slit-Robo signaling facilitates epithelial wound repair by extruding dying cells. Sci Rep. 2019;9(1):19549.

45. Berruyer C, Martin FM, Castellano R, Macone A, Malergue F, Garrido-Urbani S, et al. Vanin-1−/− mice exhibit a glutathione-mediated tissue resistance to oxidative stress. Mol Cell Biol. 2004;24(16):7214–24.

46. Ferreira DW, Naquet P, Manautou JE. Influence of Vanin-1 and Catalytic Products in Liver During Normal and Oxidative Stress Conditions. Curr Med Chem. 2015;22(20):2407–16.

47. Lebo RV, Kredich NM. Inactivation of human gamma-glutamylcysteine synthetase by cystamine. Demonstration and quantification of enzyme-ligand complexes. J Biol Chem. 1978;253(8):2615–23.

48. Cao R, Teskey G, Islamoglu H, Abrahem R, Munjal S, Gyurjian K, et al. Characterizing the Effects of Glutathione as an Immunoadjuvant in the Treatment of Tuberculosis. Antimicrob Agents Chemother. 2018;62(11).

49. Qin H, Powell-Coffman JA. The Caenorhabditis elegans aryl hydrocarbon receptor, AHR-1, regulates neuronal development. Dev Biol. 2004;270(1):64–75.

50. He G, Tsutsumi T, Zhao B, Baston DS, Zhao J, Heath-Pagliuso S, et al. Third-generation Ah receptor-responsive luciferase reporter plasmids: amplification of dioxin-responsive elements dramatically increases CALUX bioassay sensitivity and responsiveness. Toxicol Sci. 2011;123(2):511–22.

51. Dalton TP, Puga A, Shertzer HG. Induction of cellular oxidative stress by aryl hydrocarbon receptor activation. Chem Biol Interact. 2002;141(1-2):77–95.

52. Medzhitov R, Schneider DS, Soares MP. Disease tolerance as a defense strategy. Science. 2012;335(6071):936–41.

53. Naquet P, Pitari G, Dupre S, Galland F. Role of the Vnn1 pantetheinase in tissue tolerance to stress. Biochem Soc Trans. 2014;42(4):1094–100.

54. Elli L, Ciulla MM, Busca G, Roncoroni L, Maioli C, Ferrero S, et al. Beneficial effects of treatment with transglutaminase inhibitor cystamine on the severity of inflammation in a rat model of inflammatory bowel disease. Lab Invest. 2011;91(3):452–61.

55. Jeitner TM, Delikatny EJ, Ahlqvist J, Capper H, Cooper AJ. Mechanism for the inhibition of transglutaminase 2 by cystamine. Biochem Pharmacol. 2005;69(6):961–70.

56. Chen J, Lu H, Wang X, Yang J, Luo J, Wang L, et al. VNN1 contributes to the acute kidney injury-chronic kidney disease transition by promoting cellular senescence via affecting RB1 expression. FASEB J. 2022;36(9):e22472.

57. Perez-Castro L, Venkateswaran N, Garcia R, Hao YH, Lafita-Navarro MC, Kim J, et al. The AHR target gene Scinderin activates the WNT pathway by facilitating the nuclear translocation of beta-catenin. J Cell Sci. 2022.

58. Meghari S, Berruyer C, Lepidi H, Galland F, Naquet P, Mege JL. Vanin-1 controls granuloma formation and macrophage polarization in Coxiella burnetii infection. Eur J Immunol. 2007;37(1):24–32.

